# Parkinson Disease vs Aging Features Impairments in Formation and Release of a Power Grip in Real and Pantomimed String-Pulling

**DOI:** 10.64898/2026.10.29.762925

**Authors:** Maryam Kia, Surjeet Singh, Samsoon Inayat, Jon Doan, Matthew Tata, Artur Luczak, Ian Q. Whishaw

## Abstract

Parkinson disease (PD) impairs skilled reaching tasks but less is known about its effects on rhythmic bimanual actions. We used a bimanual hand-over-hand string-pulling task that requires repeated reach, grasp, pull, push and release movements. We predicted that participants with PD would show impairment in making the rotary skilled hand movement of retrieving the string, analogous to impairments observed in single handed target reaching. Nine participants with PD (mean UPDRS-III: 24.6), 16 older adults, and 10 young adults successfully performed a real string-pulling task and then pantomimed the action. Video analyses combined optical-flow estimation of whole-body movement speed, markerless 2D pose estimation of upper-limb kinematics, and expert frame-by-frame scoring of hand and finger shaping. Machine-learning classification then identified the discriminative information carried by these multilevel movement components. Participants with PD showed lower movement speed, a smaller hand workspace, and reduced hand and finger shaping, with older adults showing intermediate values. All participants grasped the real string with a power grip, but participants with PD showed reduced rotary movements of the arm and the wrists and absent of arpeggio-like sequential finger movement during grasp and release. During pantomime, participants with PD moved faster and made larger hand movements, while grasp-related hand opening and finger sequencing was reduced or absent. The findings suggest that PD is expressed across this multicomponent movement manifold, and that the movement organization changes with task demands: real string pulling requires formation and release of a power grip, whereas pantomime does not place demands on grasping ability.

**New & Noteworthy:** Parkinson disease affects multiple movement components, yet their organization during continuous action remains less studied. Using bimanual string pulling, we identified alterations particularly evident in hand opening and finger sequencing despite successful task completion. Older adults showed intermediate values on several measures, while pantomime-related changes in these measures differed across groups. These findings suggest that motor impairment in Parkinson disease includes task-dependent changes in movement organization that successful performance alone can conceal.

## Introduction

Parkinson disease (PD) is a progressive neurodegenerative disorder clinically characterized by slow movements (bradykinesia), smaller movements (hypokinesia), tremor and rigidity (Jankovic, 2008). The motor deficits are reflected in skilled upper-limb actions, including reaching, grasping, and object manipulation. In discrete, unimanual reach-to-grasp tasks, participants with PD show delayed hand shaping relative to arm transport, produce less smooth and more segmented trajectories, and show less stable spatiotemporal coordination between the transport and grasp components, particularly when precision demands increase (Castiello et al., 1993; Alberts et al., 2000; Fasano et al., 2022). Bimanual studies likewise demonstrate impaired coordination, especially when the limbs must maintain asymmetric relationships or satisfy different movement requirements (van den Berg et al., 2000; Serrien et al., 2000; Almeida et al., 2002). Extending these observations beyond isolated arm and hand movements, Whishaw et al. (2002) examined reaching for food and bringing it to the mouth as a coordinated whole-body action. Participants with PD successfully retrieved and ate the food but showed reduced forearm pronation and supination, replaced distal movements by proximal arm and trunk movements, and components normally performed concurrently became sequential. These findings suggest that the deficit is not simply a failure of movement execution but may reflect a reorganization of the motor elements used to achieve the task goal. The preservation of the task success despite changes in the contributing movement components can be considered within the central premise of the uncontrolled-manifold (UCM) framework (Scholz & Schöner, 1999). The framework proposes that the nervous system stabilizes task-relevant variables rather than prescribing a single movement solution. Because the same goal can be achieved through many combinations of joints, muscles, or finger forces, these elements can covary as a motor synergy to preserve performance despite variability in the individual components. Studies using UCM analysis have reported weaker task-stabilizing synergies and delayed or reduced preparatory changes in synergy organization before movement in PD (Park et al., 2012). Similar, though generally milder, changes have also been reported in healthy aging (Olafsdottir et al., 2007; Olafsdottir et al., 2008). Optimal feedback control (OFC) provides a complementary computational perspective in which movement being organized around task goals, with task-specific synergies emerging from the control policy (Todorov & Jordan, 2002). From this perspective changing the sensory and mechanical constraints of an action may change how its movement components are organized.

Examining how a task is accomplished, in addition to its successful completion may reveal changes in motor organization in PD. The same behavioral goal can be achieved through different combinations of body, limb, and hand movements, motivating the use of a task that allows these contributing components, and their reorganization under changing task constraints, to be examined within the same coordinated action. String pulling provides such a task (Jacobs & Osvath, 2015, Singh et al. 2019, Jordan et al., 2023). It is a continuous, rhythmic bimanual action: rather than ending after a single reach and grasp, it proceeds through repeated hand-over- hand cycles in which each hand reaches for the string, forms a grasp, pulls and pushes the string downward, releases it, and begins the next reach. The task has a long history in comparative psychology, where animals retrieve an out-of-reach reward attached to a string, allowing investigation of means–end understanding and related perceptual and cognitive processes across numerous bird and mammal species (Jacobs & Osvath, 2015). More recent studies have examined the motor organization of string pulling, including its component movements and interlimb coordination in rodents and nonhuman primates (Blackwell et al., 2018a, 2018b, 2018c; Bertrand et al., 2025). In the human version (Singh et al., 2019), participants repeatedly drew down a string suspended from an overhead pulley and then reproduced the action without the string. During real string pulling, the hands alternated, with the movement of one hand beginning before the other had completed its cycle. The fingers closed sequentially around the string to form a whole hand grasp, and participants did not visually fixate the point of contact. Without the string movements became predominantly up and down, with simpler interlimb coordination and reduced finger sequencing, suggesting that interaction with the string helps organize the repeated hand and arm movement components. We therefore used string pulling to examine how PD and aging alter the organization of movement components within the broader movement manifold. Here, the movement manifold refers to the different combinations of body, limb, and hand movements that can achieve the same task goal. We further asked how this organization changes when the participants reproduce the action without the sensory and mechanical constraints of the string. We expected PD to produce the greatest alteration in the organization of movement within this manifold, with less pronounced changes associated with aging. We also expected removal of the string to reorganize the combination of reach, grasp, used to perform the action across groups.

Participants with PD, older adults, and young adults performed string pulling with a physical string and then pantomimed the same action in its absence. We used a multilevel movement-analysis framework combining optical-flow estimation of whole-body movement, markerless 2D pose estimation of upper-limb kinematics, and expert frame-by-frame scoring of hand and finger shaping. These measures characterized whole-body movement speed, proximal arm configuration and hand transport, and distal hand opening and finger sequencing. Comparing these components across groups and task conditions allowed us to characterize change in the organization of the action. Complimentary machine-learning classification assessed which movement features contributed most to distinguishing the participant groups.

## Methods

An overview of the experimental paradigm and movement-analysis framework is provided in Figure. 1. Participants were video recorded during real string-pulling and pantomime string pulling without the string. Recordings were analyzed at three complementary levels: whole-body movement speed estimated using optical flow, proximal arm configuration and hand transport quantified using markerless 2D pose estimation, and distal hand opening and finger sequencing scored frame by frame by an experienced observer.

**Figure 1.**
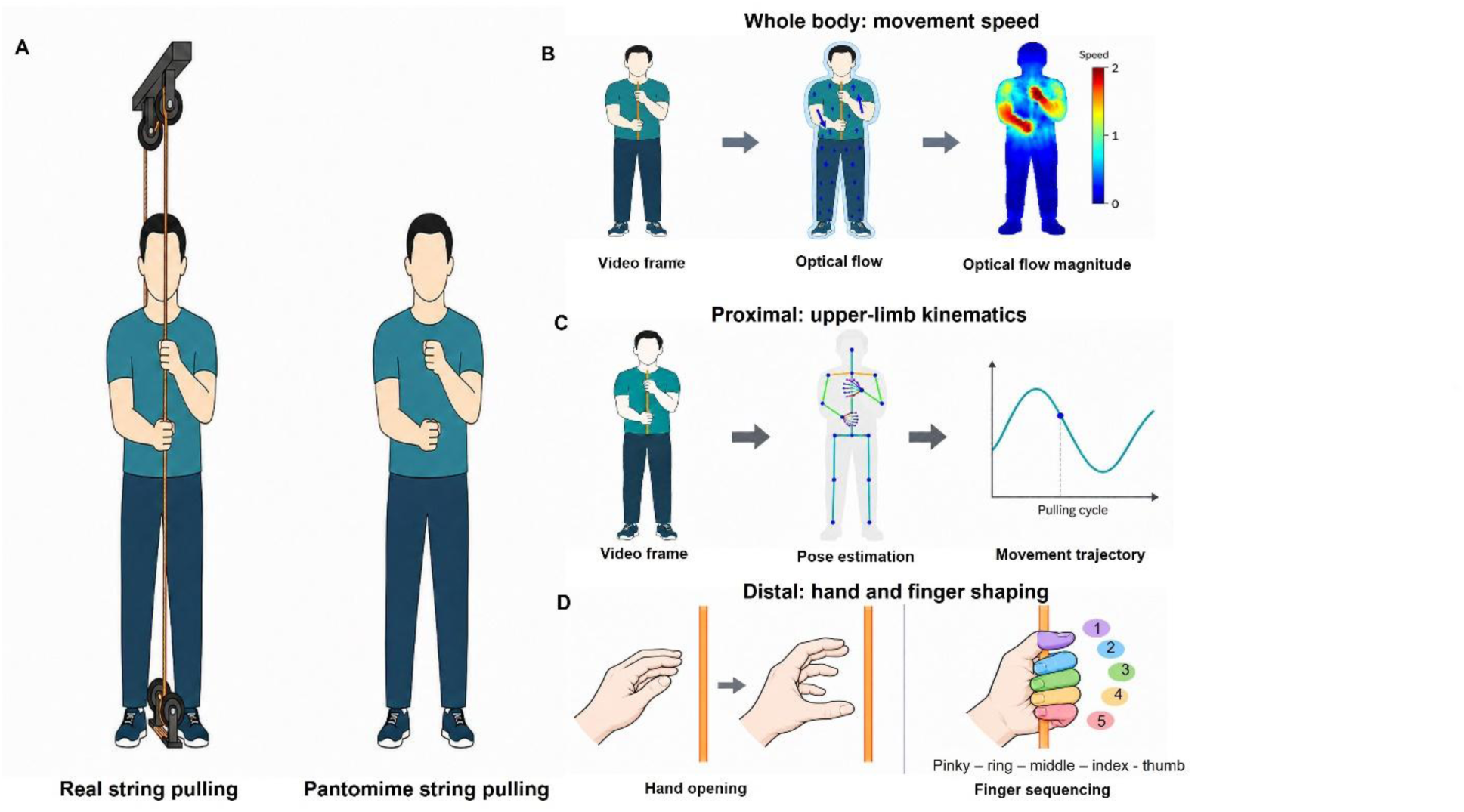
Experimental paradigm and multilevel movement-analysis framework. (A) Young adults (*n* = 10), older adults (*n* = 16), and participants with Parkinson disease (PD; *n* = 9) performed real string-pulling with the apparatus and pantomime string-pulling without the string. Video analyses quantified (B) whole-body movement speed using optical flow, (C) upper-limb kinematics using markerless two-dimensional pose estimation, and (D) hand opening and finger sequencing using observational scoring. Schematics are illustrative and do not represent participant data.

### Participants

Thirty-five participants took part: nine participants with PD, 16 age-matched older adults, and 10 young adults. Participants with PD and older adults were recruited from community exercise programs in Lethbridge, Alberta, whereas young adults were undergraduate students recruited from the University of Lethbridge. Participants with PD were tested in the on-medication state. Participants self-reported no musculoskeletal injury or other condition affecting the arm or shoulder that would prevent them from performing the string-pulling task. Participant demographic characteristics and clinical measures for the PD group are summarized in Table 1.

**Table I.** Participant demographic and clinical characteristics.

| Characteristics | Young adults | Older adults | Participants with PD |
|---|---|---|---|
| Number of participants | 10 | 16 | 9 |
| Age, years | 18–23* | 73.9 ± 5.0 | 71.0 ± 2.5 |
| Sex, male/female, n | 4/6 | 5/11 | 6/3 |
| H&Y stage | — | — | 1.5 ± 0.53 |
| S&E score, % | — | — | 87.5 ± 10.4 |
| UPDRS-III score | — | — | 24.6 ± 5.6 |

Values are means ± SD. Sex is reported as counts. \*Young adults' age is reported as a range. —, not applicable. PD, Parkinson disease; H&Y, Hoehn and Yahr; S&E, Schwab and England Activities of Daily Living; UPDRS-III, motor subsection of the Unified Parkinson Disease Rating Scale.

All participants provided written informed consent before participation, including consent for video recording and analysis. Experimental procedures were approved by the University of Lethbridge Human Subjects Research Ethics Committee and were conducted in accordance with institutional guidelines.

### String-Pulling Task

Participants performed real and pantomime string pulling adapted from the experimental paradigm described by Singh et al. (2019) (Figure 1A). The string consisted of a 7-m length of white rope suspended from the ceiling through a pulley. A knot at its end prevented the string from passing through the pulley, and an attached retrieval cord was used to return it to its starting position between trials. Participants stood facing the string and performed alternating hand-over-hand pulling movements following a verbal “go” command. In the real condition, participants pulled the suspended string until the knot reached the pulley. In the pantomime condition, the string was moved aside, and participants were asked to reproduce the same hand-over-hand movements without physically interacting with it. Each participant completed a practice trial followed by one real and one pantomime trial. Both experimental trials were video recorded and typically contained 4–7 string pulling cycles per hand.

### Movement Analysis

Figure 1B–D summarizes the three complementary levels of movement analysis: whole-body movement speed, upper-limb kinematics, and hand and finger shaping. Upper-limb kinematic measures characterized arm configuration, movement extent and workspace, timing, and speed and were grouped as proximal measures. Hand-opening and finger-sequencing scores characterized distal hand and finger shaping. The following sections describe how these measures were obtained.

### Whole-Body Movement speed

Whole-body movement speed was quantified using the optical-flow component of a MATLAB-based toolbox developed for analyzing string-pulling behavior in rodents (Inayat et al., 2020). Optical flow was computed between consecutive video frames to estimate two-dimensional motion vectors each trial. The magnitudes of these vectors were used to generate speed maps representing the spatial distribution of movement speed in each frame.

For each participant and task condition, a mean speed map was computed by averaging the speed at each pixel across trial. A region of interest (ROI) was defined by selecting pixels with values above the global mean of this map, to exclude stationary background regions and focus the analysis on areas of active movement. Whole-body movement speed was calculated as the mean speed within this ROI and expressed in cm/s, yielding one value per participant and condition.

### Upper-Limb Kinematics

Upper-limb movements were quantified using markerless two-dimensional pose estimation. Video recordings were processed using OpenPose (Cao et al., 2019), with the BODY_25 model used to estimate body keypoints and a 21-keypoint hand model used to estimate hand keypoints (Simon et al., 2017). Two-dimensional coordinates of the shoulder, elbow, wrist, and finger keypoints were extracted from each video frame. Wrist trajectories were used to characterize hand transport. Missing samples were linearly interpolated, and trajectories were smoothed using locally weighted regression (LOESS) before the analysis. Hands were classified according to wrist workspace area. The hand occupying the larger workspace was designated the dominant hand, and the other was designated the non-dominant hand.

Movement cycles were identified from the vertical wrist trajectory using automated peak-based segmentation. Upper-limb kinematic measures were organized into three categories: angular variability, movement extent and workspace, and movement timing and speed, as described below.

#### Angular Variability

Variation in upper-limb configuration was quantified using the two angles illustrated in Figure 4A. A fixed reference line was defined using the mean positions of the neck and ipsilateral shoulder across each trial. The shoulder–elbow angle described the orientation of the upper-arm segment relative to this reference whereas the shoulder–elbow–wrist angle described the elbow angle formed by the upper-arm and forearm segments. Angular variability was calculated as the standard deviation of each angular trajectory across the trial, separately for each arm and condition. Larger values indicated greater variation in arm configuration during the trial.

#### Movement Extent and Workspace

Wrist and elbow positions were measured relative to the ipsilateral shoulder to calculate the spatial measures illustrated in figure 5A. Hand height above shoulder and elbow height above shoulder were defined as the maximum vertical positions of the wrist and elbow, respectively, relative to the shoulder. Hand workspace area and elbow workspace area were calculated as the areas enclosed by the convex hulls of the corresponding two-dimensional trajectories. All measures were computed across each trial, separately for each hand and condition.

#### Movement Timing and Speed

Rise time was defined as the interval from a local minimum to the subsequent local maximum of the vertical wrist trajectory. Fall time was defined as the interval from that maximum to the subsequent minimum. These intervals primarily encompassed reaching and grasping, and pulling and release, respectively. Instantaneous hand and elbow speed were calculated as the magnitudes of the time derivatives of their respective two-dimensional wrist and elbow positions. Rise time, fall time, mean hand speed and mean elbow speed were calculated for each movement cycle and then averaged across all cycles in each trial, separately for each hand

### Hand and Finger Shaping

Hand opening and finger sequencing were assessed from video recordings using the scoring criteria described below. One experienced observer scored four movement cycles per hand in each trial.

Hand opening was assessed during lift, grasp, and release using five ordinal categories: 0, hand fully closed; 25, slight finger opening; 50, fingers half closed; 75, fingers partially flexed; and 100, hand fully open. These values represented ordered categories of hand opening rather than measured percentages. Finger sequencing was assessed during grasp and release using six scoring categories (0-5). Scores of 1–5 identified the digit initiating finger closing during grasp or finger opening during release: 1, thumb; 2, index finger; 3, middle finger; 4, ring finger; and 5, little finger. A score of 0 indicated that the hand did not open or close during the scored phase.

### Statistical and Classification Analysis

#### Statistical Analysis

Whole-body movement speed was analyzed using a two-way mixed-design ANOVA with group (young adults, older adults, and participants with PD) as the between-subject factor and task condition (real and pantomime) as the within-subject factor. Upper-limb kinematic measures were analyzed using three-way mixed design ANOVAs with group as the between-subject factor and task condition and hand as within-subject factors. Significant main effects of group were followed by Bonferroni-corrected pairwise comparisons.

Hand-opening and finger-sequencing scores were analyzed using linear mixed-effects models fitted separately for each movement phase. Fixed effects included group, task condition, hand, and all interactions with a random intercept for participant to account for repeated observations across movement cycles. Significant main effects of group were followed by Bonferroni-corrected comparisons. Statistical significance was set at α = 0.05.

### Machine-Learning Classification

Machine-learning analyses assessed whether movement measures jointly distinguished participants with PD, older adults, and young adults. For each participant, a feature vector was constructed from the behavioral measures described above, including whole-body movement speed, upper-limb kinematic measures, and hand and finger shaping measures. Each participant contributed one features vector combining whole-body movement speed, upper limb kinematic measures, and hand opening and finger sequencing scores.

#### Classification models

Two complementary classifiers were evaluated. A linear support vector machine (SVM; Cortes and Vapnik, 1995) and a gradient-boosted decision-tree classifier implemented using Extreme Gradient Boosting (XGBoost; Chen and Guestrin, 2016). The linear SVM used linear decision boundaries to distinguish participant groups, whereas XGBoost allowed nonlinear relationships and interactions among movement features. Analyses were implemented in Python using scikit-learn and the XGBoost library. Model hyperparameters and implementation details are provided in the Supplemental Material.

#### Cross-Validation and Performance Evaluation

Classification performance was evaluated using leave-one-subject-out cross-validation (LOSO-CV). In each iteration, one participant was held out for testing while the model was trained on the remaining participants. This procedure was repeated until each participant had been held out once. Performance was summarized across the pooled held out predictions using the balanced accuracy, class-specific recall, and normalized confusion matrices

#### Feature Importance

Feature importance was assessed separately for each classifier using repeated feature permutation with model retraining. Each feature was randomly shuffled across participants while the remaining features and group labels were left unchanged. The full LOSO-CV procedure, including model training, was then repeated with hyperparameters held fixed. This procedure was repeated 100 times per feature for the linear SVM and 30 times per feature for XGBoost. Feature importance was calculated as the mean decrease in balanced accuracy relative to performance with the original, unshuffled features. Importance values were interpreted in the context of the complete feature set, recognizing that correlated features may carry overlapping information. To summarize importance across the three levels of movement analysis, features were grouped into whole-body movement speed, upper-limb kinematics, and hand and finger shaping.

## Results

### Overview of the String-Pulling Movement Pattern

Figure 2 shows representative left and right hand trajectories from a young adult and a participant with PD during real and pantomime string pulling. Both participants maintained the alternating hand-over-hand pattern across repeated movement cycles in both conditions. Figures 2A-B show relatively large, regular oscillations and a broad hand workspace in the young adult. Figures 2C-D show smaller and less regular trajectories in participant with PD, particularly in real string pulling. In both participants, pantomime was accompanied by larger vertical excursions and a broader hand workspace than real string pulling. These individual movement examples illustrate the movement patterns examined in the group-level analyses analysis of whole-body movement speed, upper-limb kinematics, and hand and finger shaping below.

**Figure 2.**
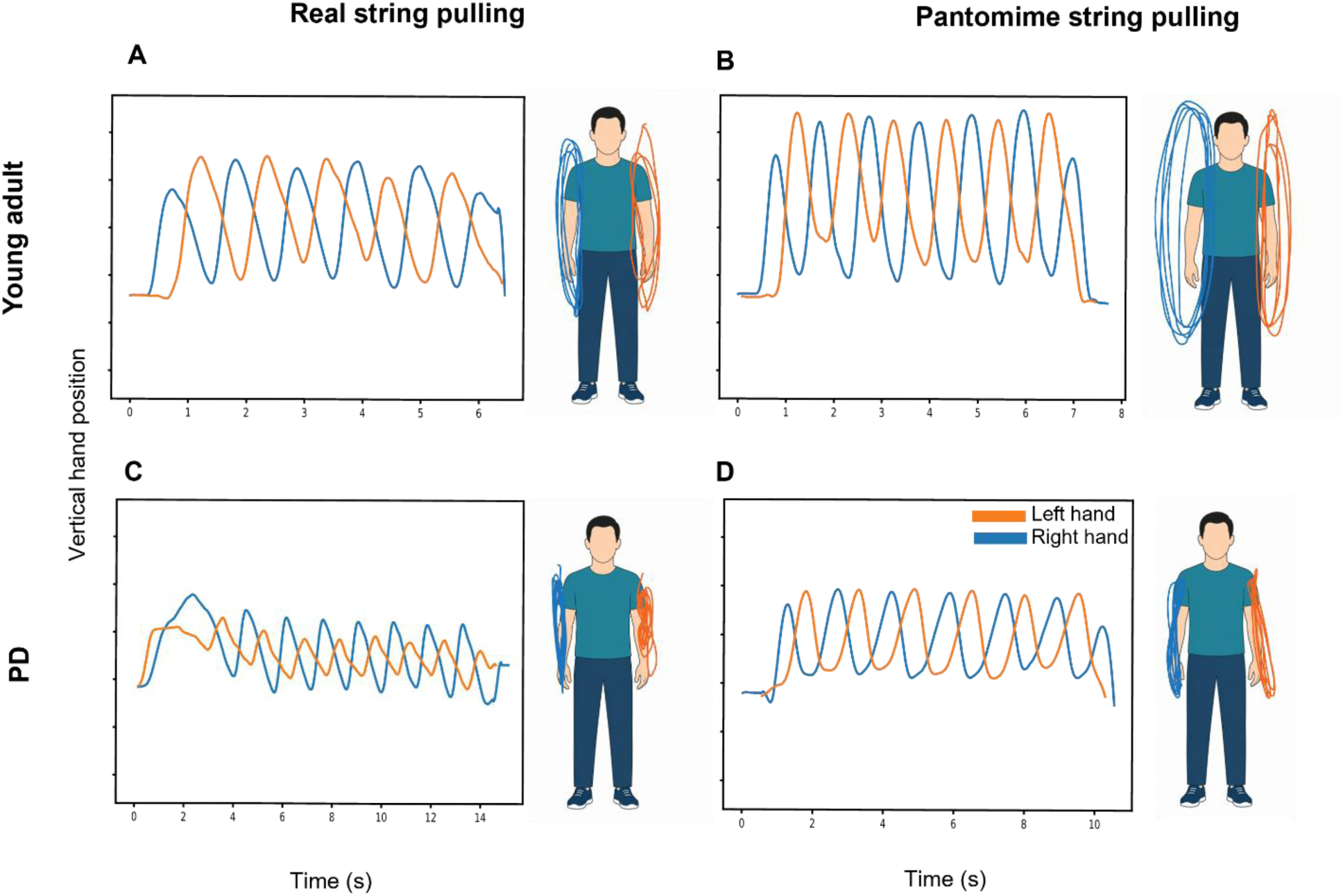
Representative hand trajectories during real and pantomime string pulling. Trajectories are shown for a young adult (A–B) and a participant with Parkinson disease (PD; C– D) during real (A, C) and pantomime (B, D) string pulling. Each panel shows vertical hand position over time alongside the corresponding spatial trajectories. Orange and blue indicate the left and right hands, respectively.

### Whole-Body Movement Speed

Figures 3A-C show representative velocity vectors, speed maps, and mean speed map. Whole-body movement speed differed across participant groups (Fig. 3D) with a significant main effect of group (F(2,32) = 5.77, p = 0.007). Bonferroni-corrected post hoc comparisons showed lower movement speed in participants with PD than in young adults (*p* = 0.006). There was also a significant main effect of task condition (F(1,32) = 6.05, p = 0.019) with higher movement speed during pantomime. The group × task interaction was not significant.

**Figure 3.**
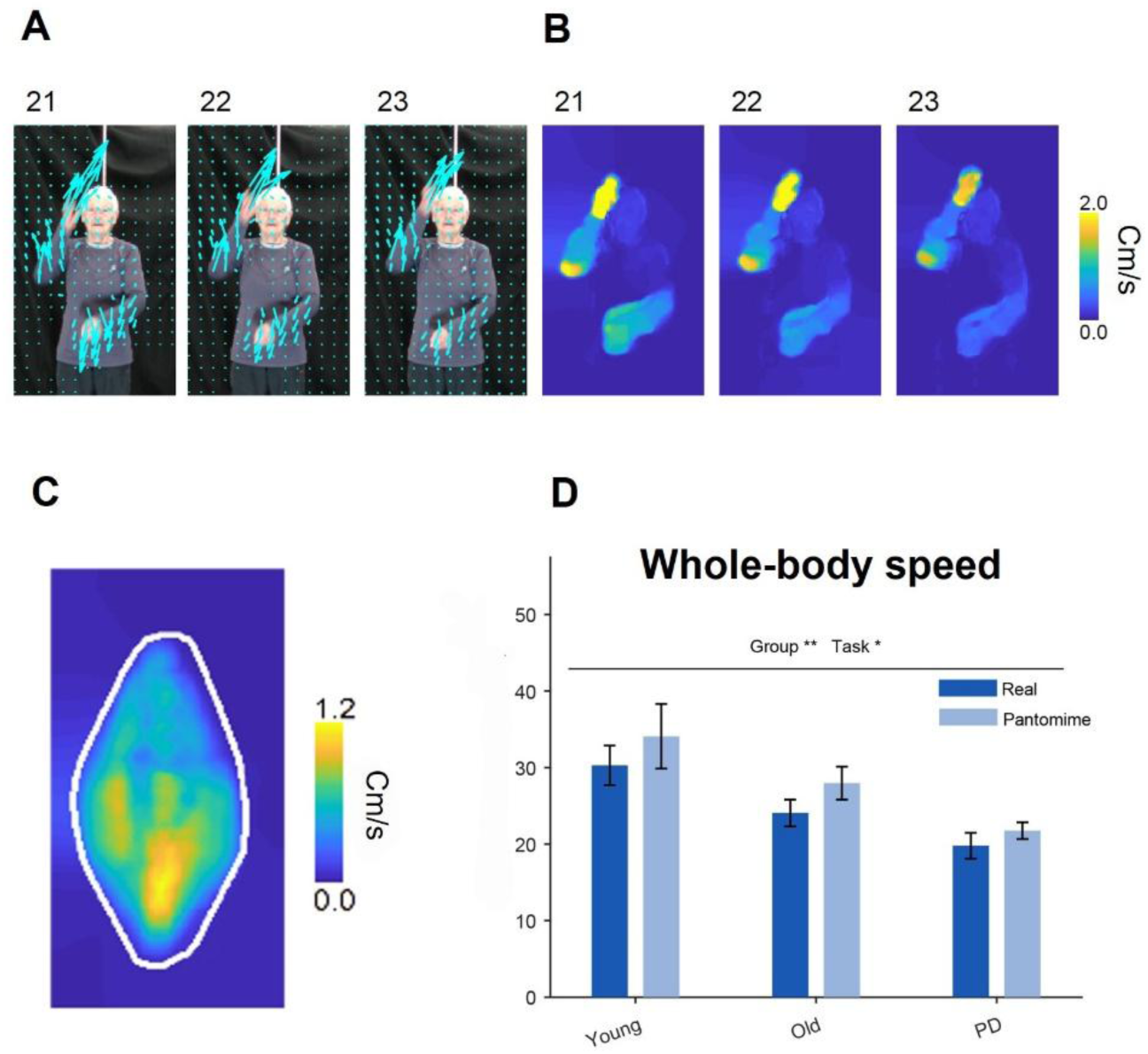
Whole-body movement speed during real and pantomime string pulling. (A) Representative optical-flow velocity vectors from three consecutive video frames illustrating the direction and magnitude of movement during string pulling. (B) Corresponding speed maps generated from the magnitude of the optical-flow vectors, with warmer colors indicating higher movement speeds. (C) Average speed map computed across the entire movement sequence. The white contour indicates the participant-specific region of interest (ROI) used to quantify whole-body movement speed. (D) Mean whole-body movement speed for young adults, older adults, and individuals with Parkinson disease (PD) during real and pantomime string pulling. Error bars represent SEM. Asterisks indicate significant main effects of group and task (\**p* < 0.05; \**p* < 0.01; \*\**p* < 0.001).

### Upper Limb Kinematics

#### Angular Variability

Figure 4 shows the shoulder–elbow (δ) and shoulder–elbow–wrist (ϕ) angles, and their variability during string pulling. Neither angular measure showed a significant main effect of group (both *p* > 0.05). Angular variability was greater during pantomime than real string pulling, with significant main effects of task condition for both the shoulder–elbow angle (F(1,32) =11.51, p =0.002; Figure 4C) and the shoulder–elbow–wrist angle (F (1,32) = 13.87, p < 0.001; Figure 4B). Shoulder–elbow–wrist angular variability was greater on the dominant than the non-dominant side (*F*(1, 32) = 4.62, *p* = 0.039), whereas shoulder–elbow angular variability showed no significant main effect of hand. No significant interaction was detected (all p > 0.05).

**Figure 4.**
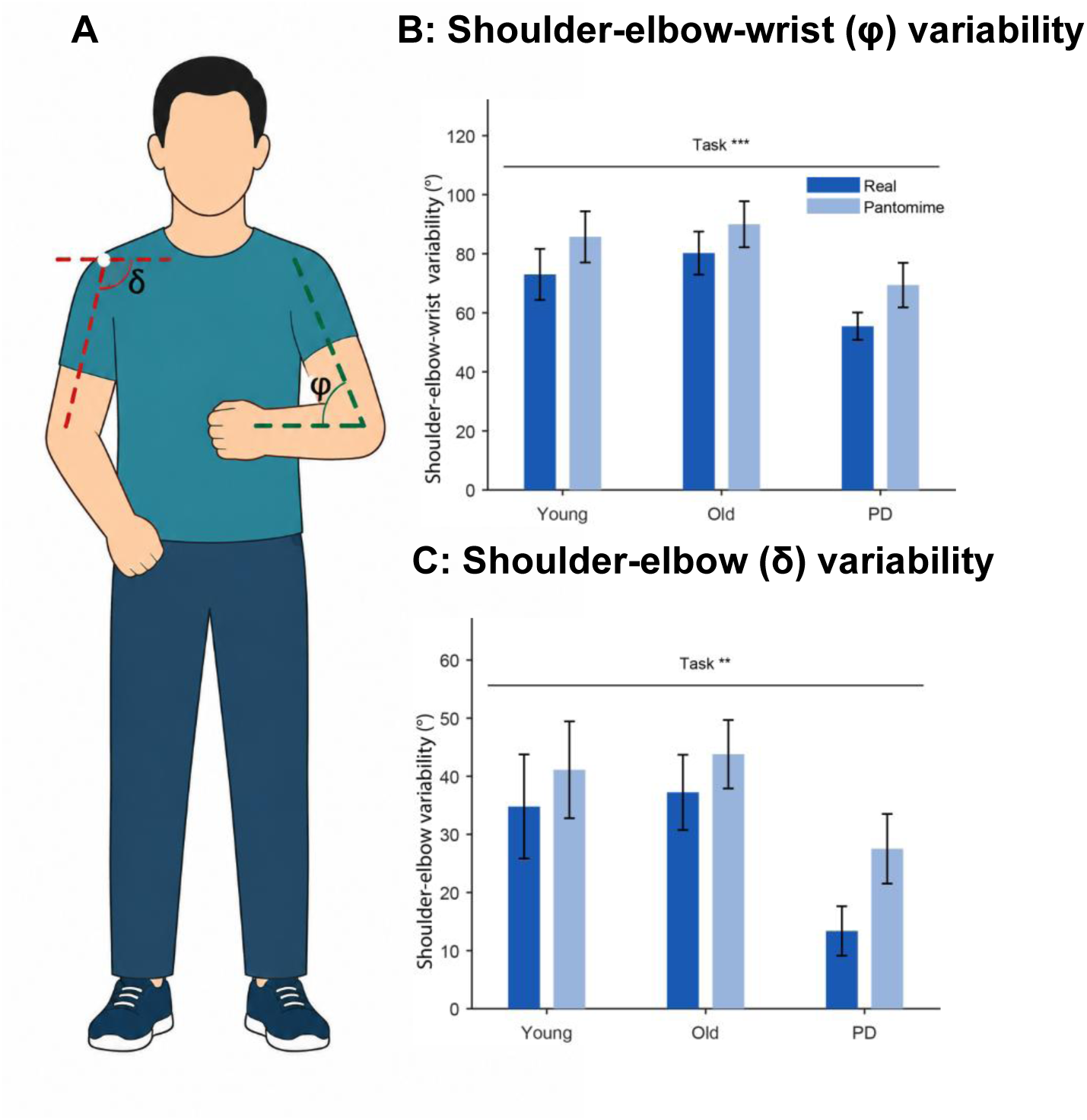
Angular variability of upper-limb joint angles during real and pantomime string pulling. (A) Definitions of the shoulder–elbow angle (δ) and shoulder–elbow–wrist angle (ϕ). Angular variability, calculated as the standard deviation across the movement sequence, is shown for (B) ϕ and (C) δ. Values were averaged across hands; bars show group means and error bars represent SEM across participants. Asterisks indicate significant main effects of group and task (\**p* < 0.05; \**p* < 0.01; \*\**p* < 0.001).

#### Movement Extent and Workspace

Figure 5 shows shoulder-referenced hand and elbow heights and their workspace areas. Significant main effects of group were found for hand height (F(2,32) = 3.43, p = 0.045; Figure 5B) and hand workspace area (*F*(2, 32) = 3.65, *p* = 0.037; Figure 5C). Bonferroni-corrected post hoc comparisons identified no significant pairwise group differences in hand height, but hand workspace area was smaller in participants with PD than in young adults (*p* = 0.048). Neither elbow height nor elbow workspace area showed a significant main effect of group (both p > 0.05).

**Figure 5.**
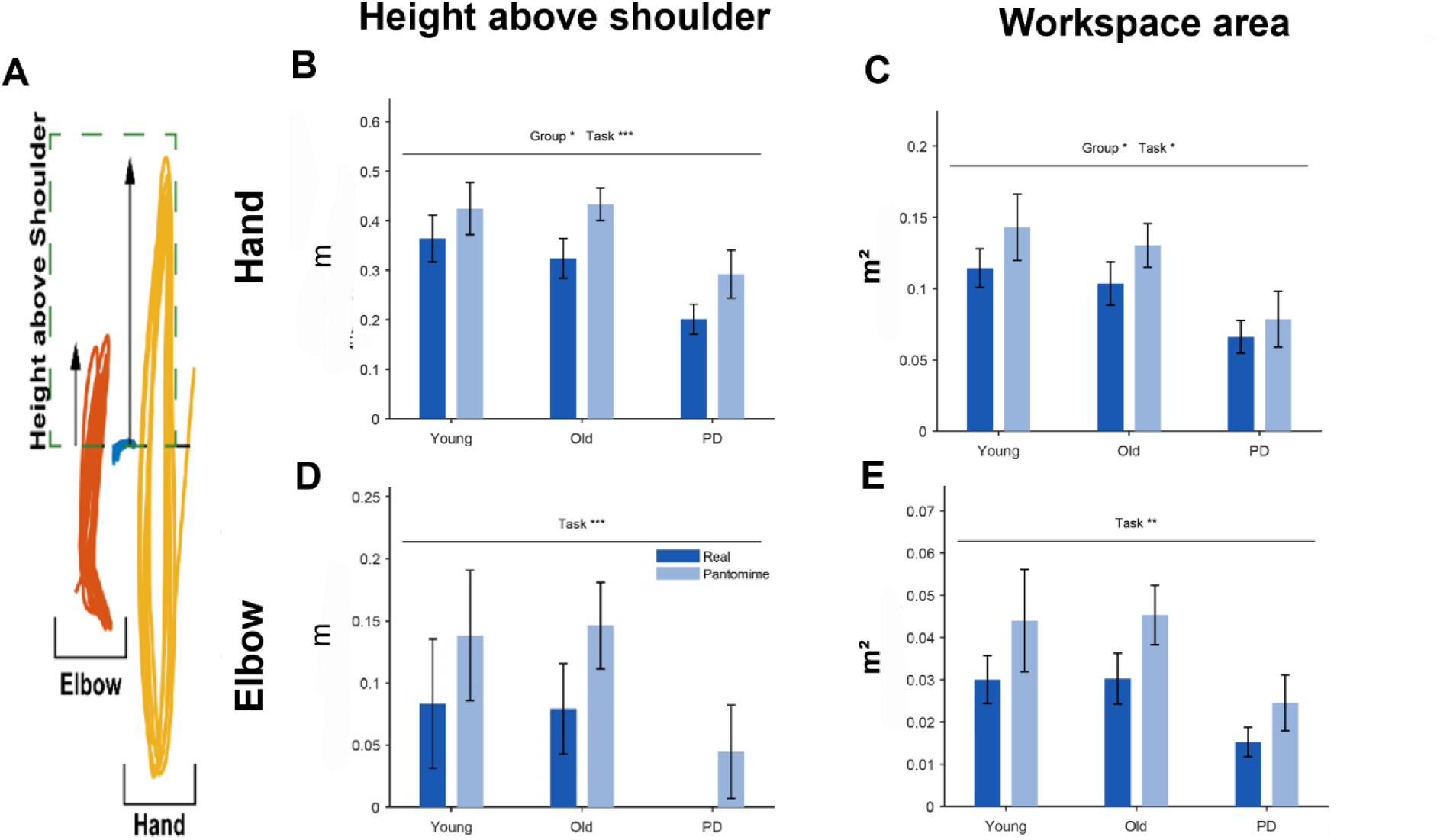
Movement extent and workspace during real and pantomime string pulling. (A) Representative hand and elbow trajectories illustrating the spatial measures. (B) Maximum hand height above the shoulder. (C) Hand workspace area. (D) Maximum elbow height above the shoulder. (E) Elbow workspace area. Heights were measured relative to the ipsilateral shoulder, and workspace areas were calculated from the convex hulls of the corresponding trajectories. Bars show group means; error bars indicate SEM. Asterisks indicate significant main effects of group and task condition (* *p* < 0.05; ** *p* < 0.01; *** *p* < 0.001).

All four measures were greater during pantomime than real string pulling. Significant main effects of task condition were observed for hand height (*F*(1, 32) = 29.90, *p* < 0.001), hand workspace area (*F*(1, 32) = 4.23, *p* = 0.048), elbow height (*F*(1, 32) = 21.25, *p* < 0.001; Figure 5D), and elbow workspace area (*F*(1, 32) = 13.07, *p* = 0.001; Figure 5E).

Significant main effects of hand were found for hand workspace area (*F*(1, 32) = 30.53, *p* < 0.001) and elbow workspace area (*F*(1, 32) = 11.25, *p* = 0.002), with both larger on the dominant side. Elbow height showed no significant main effect of hand (*p* > 0.05). No significant interactions were detected for any spatial measure (all *p* > 0.05).

#### Movement Timing and Speed

Figure 6 shows rise time, fall time, and mean hand and elbow speeds. Neither rise time nor fall time showed a significant main effect of group (both *p* > 0.05). Rise time was longer during pantomime than real string pulling (*F*(1, 32) = 30.27, *p* < 0.001; Figure 6A), whereas fall time was shorter during pantomime (F(1, 32) = 51.59, p < 0.001; Figure 6B). Neither temporal measure showed a significant main effect of hand, and no significant interactions were detected (all *p* > 0.05).

**Figure 6.**
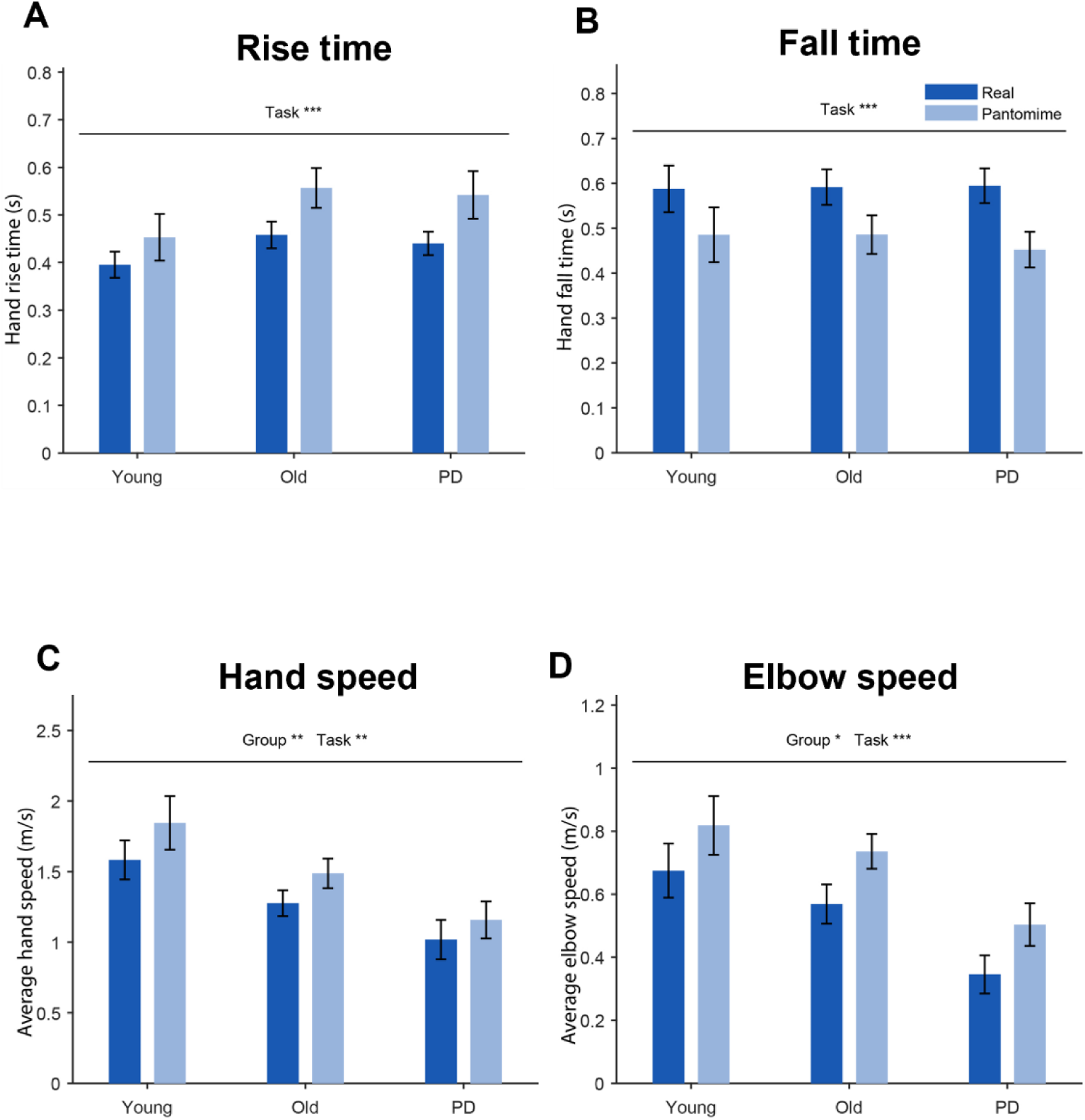
**Movement timing and speed during real and pantomime string pulling**. (A) Mean hand rise time, defined as the duration of the upward phase of the movement cycle corresponding primarily to reaching and grasping. (B) Mean hand fall time, defined as the duration of the downward phase corresponding primarily to pulling and release. (C) Mean hand speed. (D) Mean elbow speed. Data are shown for young adults, older adults, and individuals with Parkinson disease (PD) during real and pantomime string pulling. Error bars represent SEM. Asterisks indicate significant main effects of group and task (\**p* < 0.05; \**p* < 0.01; \*\**p* < 0.001).

Significant main effects of group were observed for mean hand speed (*F*(2, 32) = 5.64, *p* = 0.008; Figure 6C) and mean elbow speed (*F*(2, 32) = 5.06, *p* = 0.012; Figure 6D). Bonferroni-corrected post hoc comparisons showed lower hand and elbow speeds in participants with PD than in young adults (*p* = 0.006 and *p* = 0.012, respectively). Both measures were greater during pantomime than real string pulling, with significant main effects of task condition for hand speed (*F*(1, 32) = 13.03, *p* = 0.001) and elbow speed (*F*(1, 32) = 31.25, *p* < 0.001). Mean speeds were also higher on the dominant than the non-dominant side for both the hand (*F*(1, 32) = 12.52, *p* = 0.001) and elbow (*F*(1, 32) = 8.70, *p* = 0.006). No significant interactions were detected for either speed measure (all *p* > 0.05).

#### Hand and Finger Shaping

During real string pulling all participants grasped the string using a power grip. As previously described by Singh et al., (2019), the young adults made first contact with the string with their 5^th^ digits (pinkie) and closed the digits with an arpeggio (digits 5-4-3-2) movement in which the string was gathered into the hand with the most distal digits. At terminal grasp, the thumb sometimes played a supporting role by applying pressure to the other digits, but it was not used for grasping (Mazroui et al., 2026). When the string was released, the digits opened with an arpeggio-like sequential movement (5-4-3-2). Grip formation was similar in the older adults but digit movements, grasping and release were different as indicated by the following measures of grip formation and finger shaping.

#### Hand Opening

Figure 7 shows hand-opening scores during lift, grasp, and release phases of string pulling. Significant main effects of group were observed during lift (*F* (2, 548) = 19.86, *p* < 0.001; Figure 7A), grasp (*F*(2, 548) = 33.94, *p* < 0.001; Figure 7B), and release (*F*(2, 548) = 11.34, *p* < 0.001; Figure 7C). Bonferroni-corrected comparisons, showed higher scores in young adults than in older adults and participants with PD, and higher scores in older adults than in participants with PD. During lift, all three comparisons were significant (young versus older adults: *p* = 0.003; both comparisons involving PD: *p* < 0.001). All comparisons were also significant during grasp (all *p* < 0.001) and release (young versus older adults: *p* = 0.020; young adults versus PD: *p* < 0.001; older adults versus PD: *p* = 0.028).

**Figure 7.**
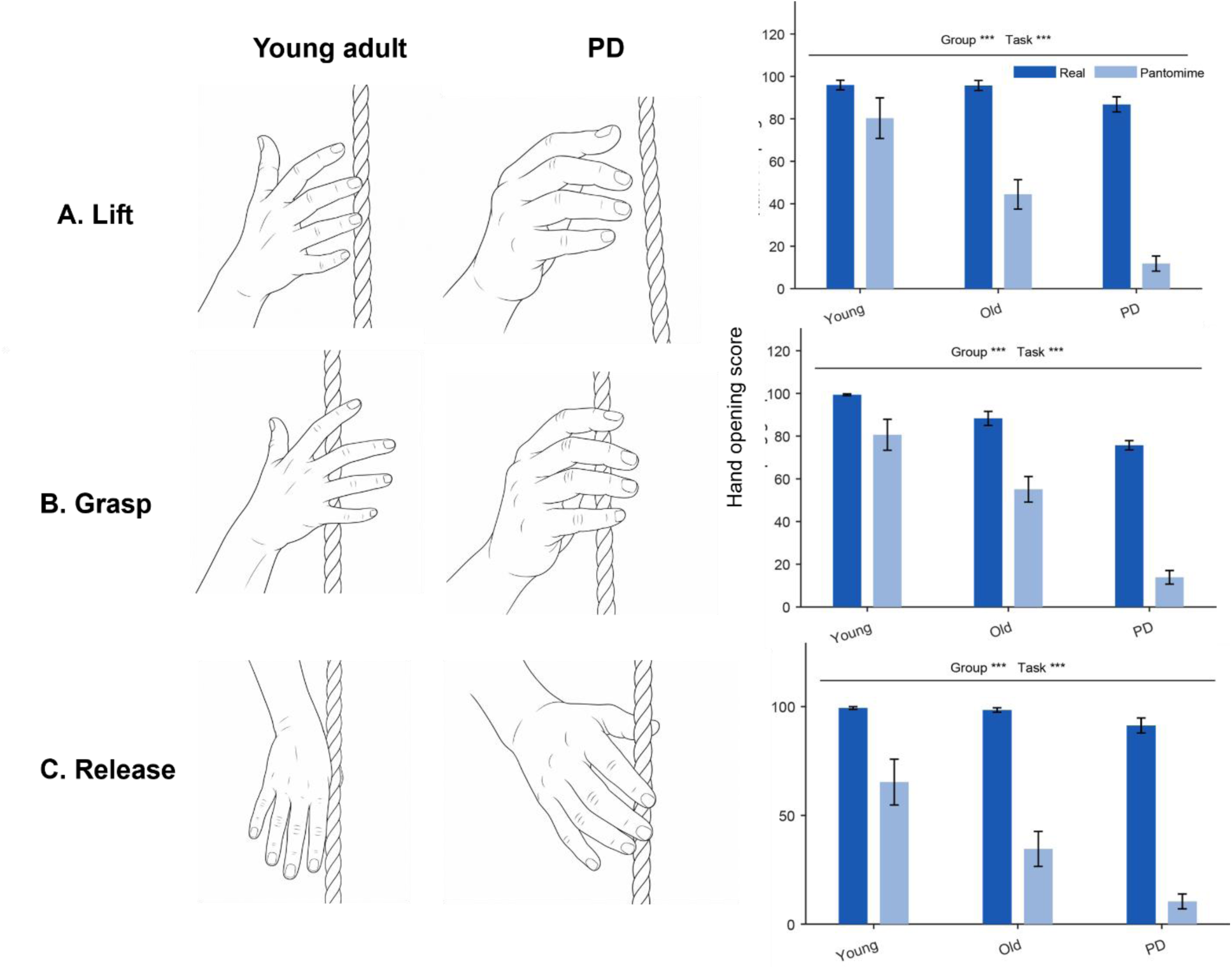
Hand opening during real and pantomime string pulling. Sketches based on selected video frames illustrate hand configurations in a young adult and a participant with Parkinson disease (PD) during real string pulling. Rows show (A) lift, (B) grasp, and (C) release, with corresponding hand-opening scores at right. Bars show group means ± SEM across participants after averaging scores across movement cycles and hands. Annotations indicate significant main effects of group and task condition (***p < 0.001).

Hand-opening scores were lower during pantomime than real string pulling, with significant main effects of task during lift (*F* (1, 548) = 799.21, *p* < 0.001), grasp (*F* (1, 548) = 562.41, *p* < 0.001), and release (*F* (1, 548) = 1232.58, *p* < 0.001). Scores were higher for the dominant than non-dominant hand during grasp (*F* (1, 548) = 5.50, *p* = 0.019) and lift (*F* (1, 548) = 4.90, *p* = 0.027), and not significant during release (*p* = 0.146).

Significant group × task interactions were observed during lift (*F* (2, 548) = 93.34, *p* < 0.001), grasp (*F* (2, 548) = 54.03, *p* < 0.001), and release (*F* (2, 548) = 57.45, *p* < 0.001). Bonferroni-corrected comparisons of task differences showed that the reduction in hand-opening scores during pantomime was greater in participants with PD than in older and young adults, and greater in older than in young adults, in all three phases (all *p* < 0.001).

A significant group × hand interaction was observed during grasp (*F*(2, 548) = 8.98, *p* < 0.001). Bonferroni-corrected comparisons showed that the dominant-minus-non-dominant difference was greater in older adults than in young adults (*p* < 0.001) and participants with PD (*p* = 0.003). During release, a significant group × task × hand interaction was observed (*F*(2, 548) = 4.45, *p* = 0.012). Follow-up comparisons showed that the difference between hands in the task-related change differed between young and older adults (*p* = 0.026) and between young adults and participants with PD (*p* = 0.029). No other interactions were significant (all *p* > 0.05).

#### Finger Sequencing

Figure 8 shows finger-sequencing scores during grasp and release. Significant main effects of group were observed during grasp (*F*(2, 548) = 8.91, *p* < 0.001) and release (*F*(2, 548) = 3.12, *p* = 0.045). Bonferroni-corrected comparisons showed lower grasp scores in older adults (*p* = 0.033) and participants with PD (*p* < 0.001) than in young adults; older adults and participants with PD did not differ significantly (*p* = 0.090). During release, scores were lower in participants with PD than in young adults (*p* = 0.039), whereas the remaining group comparisons were not significant (both *p* > 0.05).

**Figure 8.**
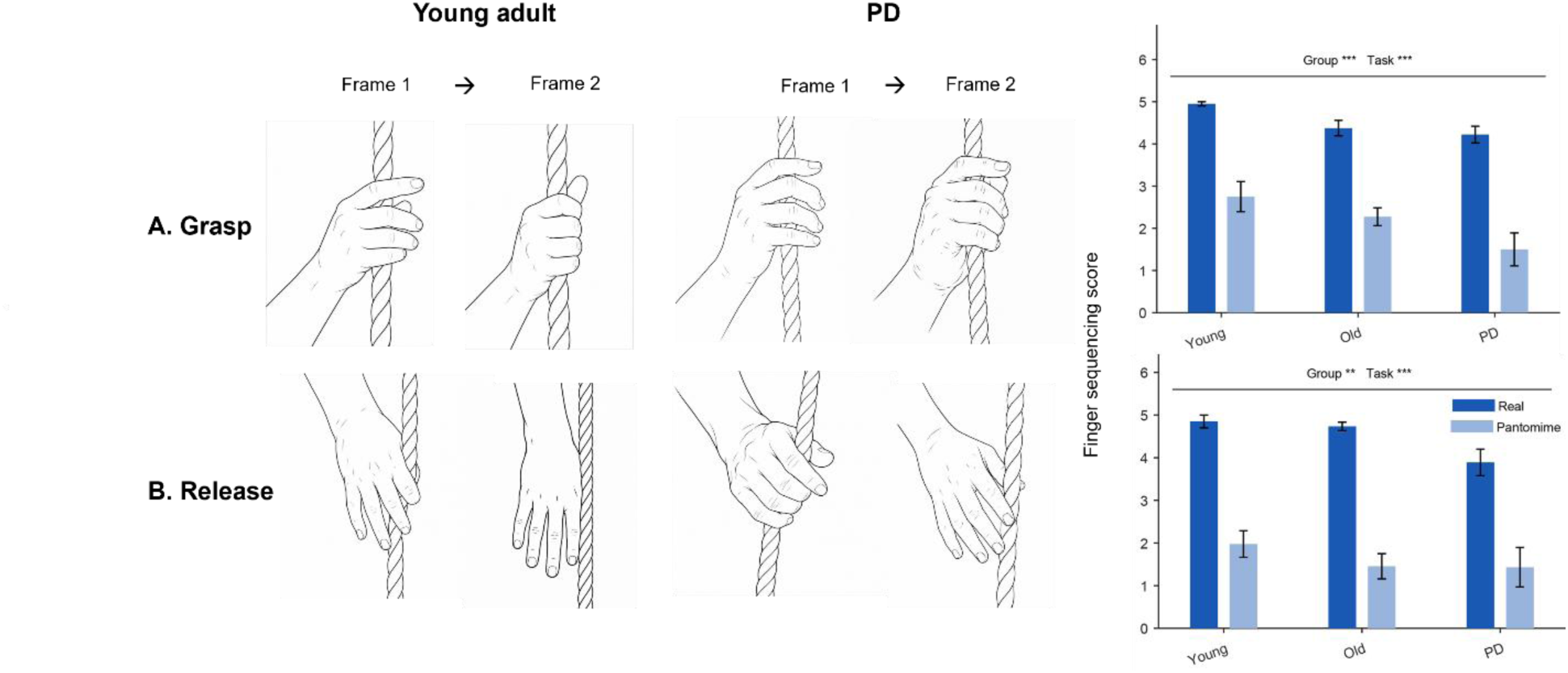
Finger sequencing during real and pantomime string pulling. Sketches based on selected video frames illustrate earlier and later hand configurations during (A) grasp and (B) release in a young adult and a participant with Parkinson disease (PD) performing real string pulling. Corresponding finger-sequencing scores are shown at right. Bars show group means ± SEM across participants after averaging scores across movement cycles and hands. Annotations indicate significant main effects of group and task condition (*p < 0.05; ***p < 0.001).

Finger-sequencing scores were lower during pantomime than real string pulling during both grasp (*F*(1, 548) = 735.20, *p* < 0.001) and release (*F*(1, 548) = 1266.75, *p* < 0.001). Scores were higher for the dominant than the non-dominant hand during grasp (*F*(1, 548) = 16.47, *p* < 0.001). The main effect of hand was not significant during release (*p* = 0.779).

Significant group × task interactions were observed during grasp (*F*(2, 548) = 4.75, *p* = 0.009) and release (*F*(2, 548) = 9.31, *p* < 0.001). Bonferroni-corrected comparisons showed a greater reduction in grasp scores during pantomime in participants with PD than in older adults (*p* = 0.008). During release, the reduction was greater in older adults than in participants with PD (*p* < 0.001). The remaining comparisons of task differences between groups were not significant.

Significant group × hand interactions were observed during grasp (*F*(2, 548) = 7.48, *p* < 0.001) and release (*F*(2, 548) = 3.87, *p* = 0.021). Bonferroni-corrected comparisons showed that the dominant-minus-non-dominant difference during grasp was greater in older adults than in young adults (*p* = 0.027) and participants with PD (*p* < 0.001). During release, this difference was greater in older than in young adults (*p* = 0.019).

An additional analysis quantified wrist-relative finger movement speed during grasp and release. Participants with PD showed lower grasp speed than young and older adults, whereas release speed was lower in both older adults and participants with PD than in young adults (Supplemental Fig. S1). Representative hand configurations and corresponding speed profiles are shown in Supplemental Fig. S2.

#### Machine-learning Classification of Participants Groups

Classification analyses combined whole-body movement speed, upper-limb kinematic measures, and hand-opening and finger-sequencing scores to distinguish young adults, older adults, and participants with PD.

#### Linear Support Vector Machine

The linear SVM achieved a balanced accuracy of 0.67 under LOSO-CV. Recall was 0.70 for young adults, 0.44 for older adults, and 0.88 for participants with PD (Fig. 9A). Of the 13 misclassifications, nine occurred between young and older adults and four occurred between older adults and participants with PD. No misclassifications occurred between young adults and participants with PD

**Figure 9.**
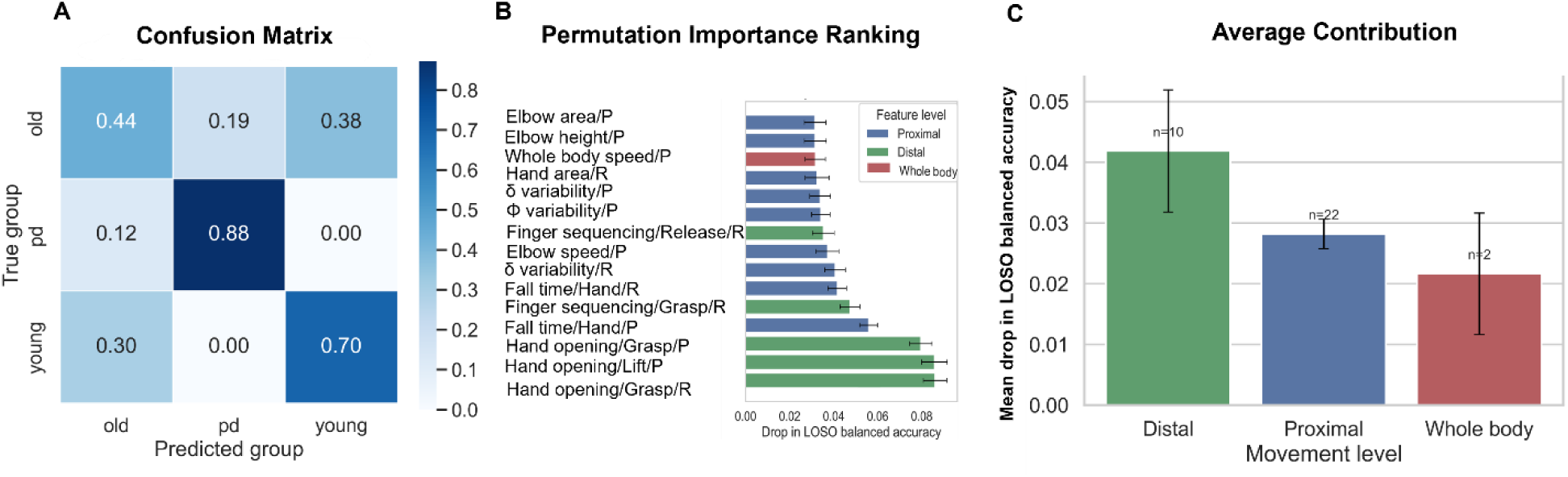
Linear SVM classification of participant groups using multi-level movement features. (A) Confusion matrix showing leave-one-subject-out (LOSO) classification performance for the linear support vector machine (SVM). Values represent recall normalized within each true group. (B) Permutation-importance analysis showing the individual features producing the largest decreases in balanced accuracy when permuted. Features are color-coded according to movement level (distal, proximal, whole-body). Error bars represent mean ± SEM across LOSO folds. (C) Average permutation importance grouped by movement level. Distal features exhibited the largest overall contribution to classification performance, followed by proximal and whole-body measures. Numbers above bars indicate the number of features within each category.

Hand-opening measures occupied the three highest positions in the permutation-importance ranking (Fig. 9B). Hand opening during real string-pulling grasp produced the largest mean decrease in balanced accuracy when shuffled and the classification procedure repeated. Other highly ranked features included fall time during pantomime and finger sequencing during real string-pulling grasp.

Mean importance per feature was highest for hand-opening and finger-sequencing measures (0.042), followed by upper-limb kinematic measures (0.028) and whole-body movement speed (0.022; Fig. 9C).

#### Extreme Gradient Boosting

XGBoost achieved a balanced accuracy of 0.83 under LOSO-CV, compared with 0.67 for the linear SVM. Recall was 0.80 for young adults, 0.81 for older adults, and 0.88 for participants with PD (Fig. 10A). Of the six misclassifications, three occurred between young and older adults, two between older adults and participants with PD, and one involved a young adult classified as having PD.

**Figure 10.**
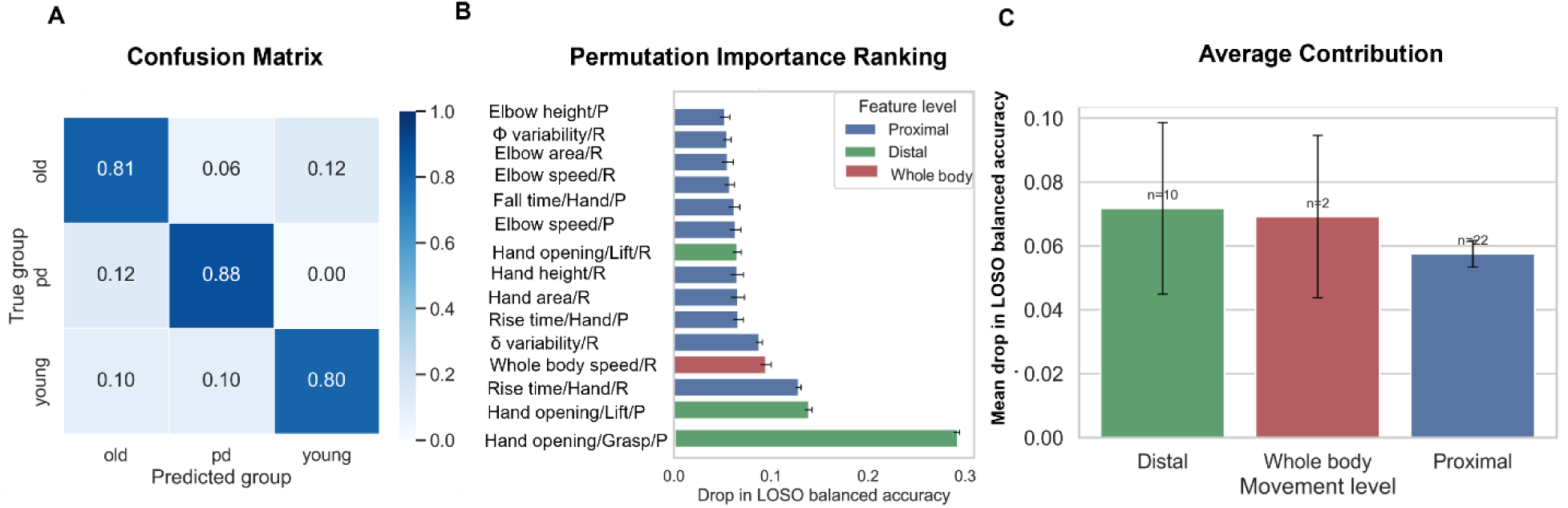
XGBoost classification of participant groups using multi-level movement features. (A) Confusion matrix showing LOSO classification performance for the XGBoost classifier. Values represent recall normalized within each true group. (B) Permutation-importance analysis showing the individual features associated with the largest decreases in balanced accuracy when permuted. Features are color-coded according to movement level (distal, proximal, whole-body). Error bars represent mean ± SEM across LOSO folds. (C) Average permutation importance grouped by movement level. Distal features exhibited the largest average contribution to classification performance, followed by whole-body and proximal measures. Numbers above bars indicate the number of features within each category.

Hand opening during pantomime grasp had the highest permutation importance, with a mean decrease in balanced accuracy of 0.291 (Fig. 10B). A second pantomime hand-opening measure ranked next, followed by rise time and whole-body movement speed during real string pulling. Mean importance per feature was 0.072 for hand-opening and finger-sequencing measures, 0.069 for whole-body movement speed, and 0.058 for upper-limb kinematic measures (Fig. 10C).

## Discussion

Participants with PD were compared with older and young adults during real and pantomime hand over hand string pulling. The PD group successfully pulled the string and completed repeated alternating cycles, but showed changes in several components of the action, including reaching and grasping. The most pronounced group differences occurred in hand closing and opening with the absence of arpeggio-like finger movements during formation and release of the power grip in participants with PD. Removing the string produced a reorganization of the action: grasp-related hand opening and finger sequencing diminished, whereas upper-limb movements for reaching became larger and faster. These findings suggest PD simplifies the organization of the movement manifold despite preserved task completion.

PD affected multiple components of string pulling despite successful task completion. Participants with PD moved more slowly, showed lower hand and elbow speeds, and a smaller hand workspace, alongside differences in hand and finger shaping. These findings are consistent with previous descriptions of impaired skilled movements in PD (Whishaw et al., 2002; Fasano et al., 2022), while extending the observation of reduced rotary movements during skilled reaching to a continuous bimanual task requiring repeated formation and release of a power grip. Older adults showed intermediate values on several measures, suggesting that aging and PD affect related dimensions of the action but to different degrees. These movement differences were not apparent from task completion alone: participants with PD advanced the string while maintaining the repeated hand-over-hand sequence. These findings highlight two complementary aspects of motor performance: whether the task goal is achieved and how the action is organized to achieve it. Assessing task completion alone would therefore overlook the movement alterations.

Participants used a power grip to grasp the string, and the largest group differences concerned the organization of grasp and release. In young adults, the fingers closed around the string in an arpeggio-like sequence progressing from the little finger to the index finger (digits 5-4-3-2), with a similar opening sequence during release. During real string pulling, participants with PD showed less distinct finger sequencing: the digits tended to open and close together, and the hand remained in a relatively horizontal orientation. During pantomime, they reproduced string pulling with the fingers largely closed. The reduced rotary hand movements during real string pulling resemble those previously described during precision grasping of food items (Whishaw et al., 2002), extending this observation to a task requiring repeated formation and release of a power grip. This extension is notable because power and precision grips differ in their neural demands. Power grip is described as a phylogenetically older form of prehension in which the digits act together to enclose an object against the palm, whereas precision grip requires greater fractionation and independent control of the digits (Napier, 1956; Lemon, 1997, 2008). The evolution of dexterous precision grip in primates is closely associated with direct cortico-motoneuronal projections from motor cortex to spinal motoneurons controlling the hand, which contribute to selective activation of individual digit muscles (Lemon, 2008; Muir & Lemon, 1983; Bennett & Lemon, 1996). More broadly distributed descending pathways, including reticulospinal projections, can contribute to coordinating groups of hand muscles during whole-hand actions. The impairment observed here therefore suggests that altered hand control in PD extends beyond the highly fractionated movements required for precision grip to the more coordinated organization of a power grip. The present observations highlight that although the string is held with a power grip, the arpeggio-like sequence gives individual fingers distinct roles in forming and releasing the grasp. In participants with PD, this organization appeared simplified, with the fingers tending to move together rather than contributing sequentially. The reduced hand opening during pantomime may also relate to the tendency toward flexed hand postures described when at rest in some individuals with PD, including striatal hand deformities and dystonic clenched-fist configurations (Ashour et al., 2005; Jankovic, 2008; Brogren et al., 2022). Such postures may become more apparent when the external requirement to open the hand and shape the digits around an object is removed.

During string pulling, each reach transports the hand upward toward the string and positions it for the next grasp, while alternating arm movements maintain the continuous hand-over-hand action. In young adults, these reaches formed relatively regular alternating excursions with a broad spatial extent. Participants with PD maintained the alternating pattern but showed a smaller hand workspace and lower hand and elbow speeds than young adults. These findings are consistent with previous reports of slower upper-limb movements and reduced elbow flexion– extension range during a hand-to-mouth task in PD (Corona et al., 2018). The smaller movement extent also resembles the reduced stride length observed in Parkinsonian gait, where forward progression can be maintained through smaller steps (Morris et al., 1994). String pulling provides a continuous bimanual setting in which to examine movement extent across successive reach–grasp cycles. When the string was removed, this pattern changed: reaching movements became larger and faster, with greater hand and elbow excursion and increased upper-limb joint variability. One possible explanation for the smaller hand workspace in PD during real string pulling is that reaching is constrained by difficulties in grasp formation, including reduced rotary hand movements. To grasp the string, upward hand transport must be coordinated with hand rotation, opening, and sequential finger recruitment. When these distal movements are diminished the reach may remain spatially constrained so that the string can still be contacted and grasped. Unlike discrete reaching tasks with a fixed target location, string pulling allows greater freedom in determining the spatial extent of each repeated reach. The smaller workspace in PD may therefore reflect a movement solution shaped partly by the demands of grasping the string; when those demands are removed, the reach expands.

The present findings can be interpreted within the frameworks of motor abundance and optimal feedback control. The UCM framework proposes that task-relevant outcomes are stabilized through covariation among contributing motor elements, allowing variation in their individual contributions while preserving the overall outcome (Scholz & Schöner, 1999; Latash, 2024).

Formal UCM studies have shown that PD is associated with weaker task-stabilizing covariation among motor elements and motor synergies, as well as reduced or delayed anticipatory adjustments of those synergies (Vaz et al., 2019). Previous studies have reported that individuals with PD can produce accurate task-level forces despite weaker multifinger synergies and impaired anticipatory synergy adjustments (Park et al., 2012; Vaz et al., 2019). Aging has also been associated with reduced bilateral synergies without an obvious failure of task performance (Kang et al., 2019). In the present study, participants with PD preserved the global goal of advancing the string despite alterations in whole-body, proximal, and distal movement components, suggesting that the task goal remained within the achievable solution manifold, but the combination of motor elements used to reach that goal was shifted. This is particularly relevant to prehension, in which a proximal transport component brings the hand toward the object while a distal grasp component shapes the hand in anticipation of contact (Jeannerod, 1984, 1986).

In the present study, the reduction in hand opening and finger sequencing in PD indicates that successful string pulling was achieved despite alteration of the distal grasp component, consistent with previous reports of impaired hand shaping, finger individuation, unintended digit coupling, and multifinger synergy organization in PD (Castiello et al., 1993; Agostino et al., 2003; Park et al., 2012). Optimal feedback control (OFC) provides a complementary perspective on how such reorganization may occur. In this framework, movement solutions emerge from task-specific goals, sensory feedback, and costs rather than from a fixed trajectory (Todorov & Jordan, 2002; Kelley & Kauffman, 2020). During real string pulling, reaching, hand shaping, rotary movements, grasping, and pulling are mechanically coupled; thus, a smaller-amplitude reach in PD may represent a task-dependent solution that reduces the demands placed on impaired distal components while still allowing the string to be captured and advanced. When the string was removed, these object-related constraints no longer had to be satisfied: reaching became larger, faster, and more variable, whereas grasp-related hand opening and finger sequencing diminished. This pattern is consistent with evidence that reach and grasp are reorganized when object-related information is removed or when the requirements for contacting and securing an object change (Kuntz & Whishaw, 2016; Mazrouei et al., 2026). This pattern suggests that removal of the string changed the set of task-relevant constraints, allowing proximal movement dimensions to vary more freely while eliminating the sensory and mechanical events that normally organize the distal grasp. The expansion of reach amplitude during pantomime further argues against a fixed inability of participants with PD to generate larger movements and instead suggests that the reduced amplitude observed during real string pulling emerged as part of a task-specific motor solution (Mazzoni et al., 2007; Baraduc et al., 2013).

Several limitations should be considered. Participants with PD were assessed in their usual medicated state, which reflects everyday functioning but may have reduced the magnitude of motor deficits observed in the present study, as dopaminergic medication improves motor symptoms and aspects of movement coordination in PD (Park et al., 2012; Alaei et al., 2025). Future studies comparing medicated and unmedicated performance could clarify the contribution of dopaminergic mechanisms to movement organization during continuous action. The PD cohort consisted primarily of individuals with mild to moderate impairment, limiting conclusions regarding later disease stages. Examination of a broader spectrum of disease severity would help determine how proximal and distal movement deficits evolve with disease progression.

Participants completed one trial per condition, with multiple movement cycles within each trial (4–7 per hand). This design provided several within-trial movement samples while minimizing potential fatigue and repetition-related changes in performance both of which can influence motor behavior and coordination (Lou et al., 2003; Vaz et al., 2019; Khalil et al., 2024). Future studies with additional trials could further characterize within-subject variability and adaptation across repetitions. Finally, the present findings suggest that string pulling may have value as a behavioral assay for characterizing movement organization in Parkinson disease. Future work combining quantitative movement analysis with neural or clinical measures may help determine whether alterations in grasp-related control provide useful markers of disease progression or treatment response.

In conclusion, continuous bimanual string pulling reveals that upper-limb impairment in PD is not confined to reduced movement but reflects a multilevel reorganization of the action. Although impairments were expressed across movement components, they were most pronounced in the distal organization of grasp and release, where the sequential shaping of the hand was diminished. Removing the string further altered how these components were organized, demonstrating that PD affects not only the execution of individual movement components, but how those components are assembled into a coordinated action. Viewing motor impairment at this level may advance the study of motor control in Parkinson disease and inform rehabilitation approaches that target the coordination of functionally integrated actions rather than isolated movement components.

## Supporting information

Suppplemental Material

## Supplemental Material

Supplemental material includes methods and results for the analysis of finger movement speed during grasp and release, group comparisons (Supplemental Fig. S1), and representative hand configurations and speed profiles (Supplemental Fig. S2). The supplemental file is provided as a separate file accompanying this submission

## Data Availability

Analysis code and deidentified numerical data will be made available at https://github.com/niktaaan/string-pulling-parkinsons-aging upon publication.

## Author Contributions

S.S., J.D., and I.Q.W. conceived and designed research; S.S. and J.D. performed experiments; M.K., S.S., S.I., and I.Q.W. analyzed data; M.K. prepared figures; M.K. and I.Q.W. drafted manuscript; M.K., S.S., S.I., M.T., A.L., and I.Q.W. edited and revised manuscript. All authors approved the final version of the manuscript.

