## Supplementary material for "Parkinson Disease vs Aging Features Impairments in Formation and Release of a Power Grip in Real and Pantomimed String-Pulling": Suppplemental Material

### S1. Finger movement speed during grasp and release

#### Methods

An additional video-based analysis quantified finger movement speed during grasp and release to complement the observational measures of hand opening and finger sequencing. Hand-keypoint trajectories were extracted using the OpenPose 21-keypoint hand model. Missing coordinates were linearly interpolated, and trajectories were smoothed using local regression. Hand keypoints were expressed relative to the wrist position in each frame to remove wrist translation. Optical flow was calculated from rendered sequences of the wrist-centered hand skeleton. The magnitudes of the resulting motion vectors provided an estimate of finger movement speed relative to the wrist. Grasp and release events were identified at the upper and lower turning points of the vertical wrist trajectory, respectively. Mean speed was calculated over nine-frame windows centered on each event and summarized for each participant, task condition, and hand. Grasp and release speeds were analyzed separately using three-way mixed-design ANOVAs, with group (young adults, older adults, and participants with Parkinson disease [PD]) as the between-subject factor and task condition (real, pantomime) and hand (dominant, non-dominant) as within-subject factors. Pairwise group comparisons were Bonferroni-corrected. Statistical significance was set at $p < 0.05$

#### Results

Finger movement speed during grasp differed across groups ($F(2, 32) = 6.30$, $p = 0.005$; Supplemental Fig. S1A). Participants with PD showed lower grasp speed than young adults (p =

0.008) and older adults (p = 0.013), whereas young and older adults did not differ significantly (p = 1.000). Neither task condition nor hand had a significant main effect, and no interactions were significant. Finger movement speed during release also differed across groups (F(2, 32) = 10.18, p < 0.001; Supplemental Fig. S1B). Young adults showed higher release speed than older adults (p = 0.034) and participants with PD (p < 0.001). Older adults and participants with PD did not differ significantly (p = 0.065). Release speed was higher during real than pantomime string pulling (F(1, 32) = 22.11, p < 0.001). The main effect of hand and all interactions were nonsignificant. Supplemental Figure S2 illustrates changes in hand configuration and the corresponding wrist-relative finger movement speed profiles across the grasp and release windows. In the illustrated real-release profile, speed was highest early in the window and declined toward its end. The grasp profiles showed smaller fluctuations across the window. These profiles illustrate the time course underlying the event-averaged measures in Supplemental Fig. S1.

**Supplemental Figures**

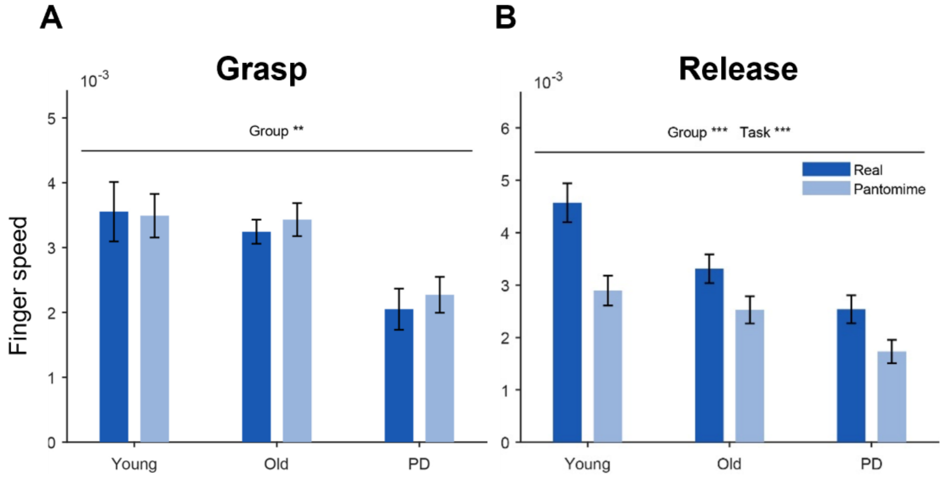


**Supplemental Figure S1. Finger movement speed during grasp and release.** Mean wrist-relative finger movement speed during (A) grasp and (B) release in young adults, older adults, and participants with PD during real and pantomime string pulling. Values were averaged across dominant and non-dominant hands within each participant and task condition. Bars show group means; error bars indicate

SEM across participant means. Annotations indicate significant omnibus main effects of group and task condition (**p < 0.01; ***p < 0.001).

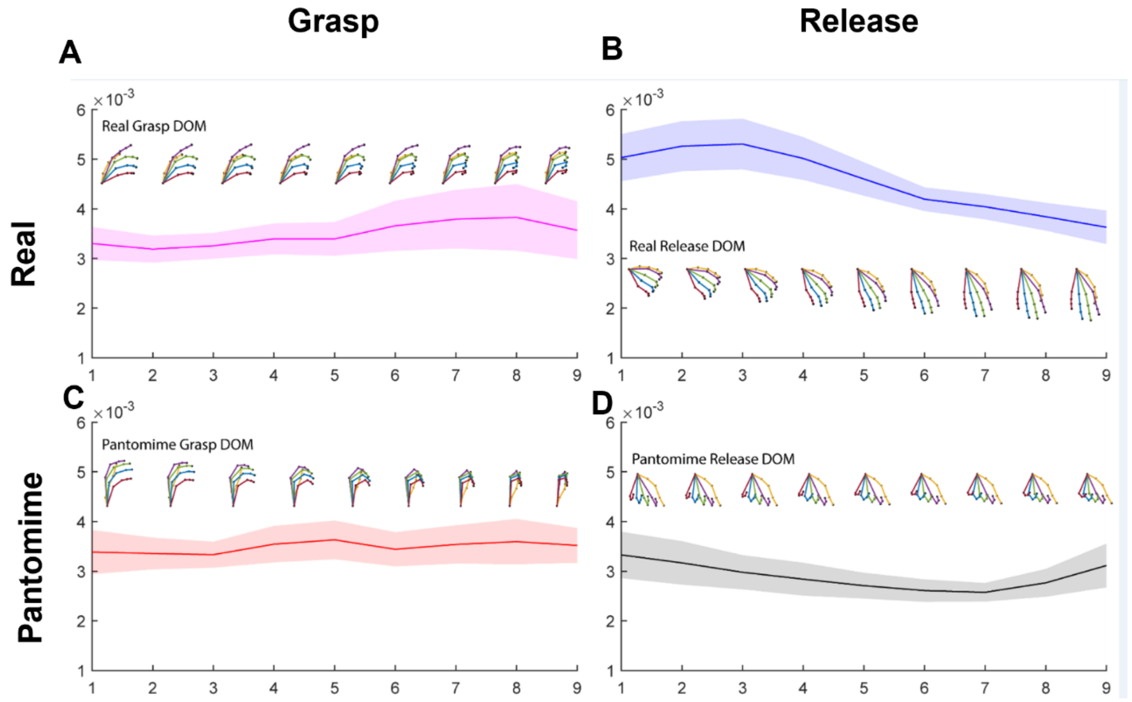


**Supplemental Figure S2. Hand configurations and wrist-relative finger movement speed during grasp and release in young adults.** Dominant-hand movements are shown during (A) real grasp, (B) real release, (C) pantomime grasp, and (D) pantomime release. Each panel spans a nine-frame window centered on the identified grasp or release event. Hand skeletons illustrate successive configurations within the window; their placement above or below the curves is for visualization and does not represent speed. Curves show mean finger movement speed estimated from optical flow applied to rendered wrist-centered hand skeletons. Shaded bands indicate SD across participants. Horizontal-axis values indicate frame number within the window. DOM, dominant hand.

### S2. Machine-learning implementation details

#### Linear support vector machine.

A linear support vector machine was implemented using scikit-learn's SVC. Features were standardized using the mean and standard deviation of the training participants within each leave-one-subject-out cross-validation (LOSO-CV) fold, and the same transformation was applied to the held-out participant. The regularization parameter was fixed at $C = 0.1$ following preliminary model evaluation. Class weighting was set to “balanced” to account for unequal group sizes.

**Extreme gradient boosting (XGBoost).**

Hyperparameters were selected using a constrained grid search evaluated with LOSO-CV. The search included the number of estimators (50, 100), maximum tree depth (2, 3), learning rate (0.03, 0.1), subsampling fraction (0.8, 1.0), and column subsampling fraction (0.8, 1.0). The combination yielding the highest balanced accuracy was selected: 50 estimators, a maximum tree depth of 2, a learning rate of 0.1, a subsampling fraction of 1.0, and a column subsampling fraction of 0.8.
